# JNK signaling controls branching, nucleokinesis, and positioning of centrosomes and primary cilia in migrating cortical interneurons

**DOI:** 10.1101/2020.01.30.927855

**Authors:** Skye E. Smith, Nicholas K. Coker, Eric S. Tucker

## Abstract

Aberrant migration of inhibitory interneurons can alter the formation of cortical circuitry and lead to severe neurological disorders including epilepsy, autism, and schizophrenia. However, mechanisms involved in directing the migration of these cells remain incompletely understood. In the current study, we used live-cell confocal microscopy to explore the mechanisms by which the c-Jun NH_2_-terminal kinase (JNK) pathway coordinates leading process branching and nucleokinesis, two cell biological processes that are essential for the guided migration of cortical interneurons. Pharmacological inhibition of JNK signaling disrupts the kinetics of leading process branching, rate and amplitude of nucleokinesis, and leads to the rearward mislocalization of the centrosome and primary cilium to the trailing process. Genetic loss of *Jnk* from interneurons corroborates our pharmacological observations and suggests that important mechanics of interneuron migration depend on the intrinsic activity of JNK. These findings suggest that JNK signaling regulates leading process branching, nucleokinesis, and the trafficking of centrosomes and cilia during interneuron migration, and further implicates JNK signaling as an important mediator of cortical development.

**Summary Statement:** Loss of JNK signaling reduces growth cone branching frequency, limits interstitial side branch duration, alters rate and amplitude of nucleokinesis, and mislocalizes centrosomes and primary cilia in migrating cortical interneurons.

## INTRODUCTION

During embryonic development, cortical interneurons are born in the medial and caudal ganglionic eminences (MGE and CGE) of the ventral forebrain and then migrate long distances to reach the place of their terminal differentiation in the overlying cerebral cortex (Miyoshi et al., 2010; Nery et al., 2002; Wichterle et al., 1999; Xu et al., 2004). While navigating their environments, cortical interneurons must integrate extracellular guidance cues with intracellular machinery in order to reach the cortex, assemble and travel in tangentially oriented streams, and disembark from streams at the correct time and place to properly infiltrate the cortical plate. Two cellular mechanisms that enable interneurons to make these complex migratory decisions are leading process branching, where cortical interneurons dynamically remodel their leading processes to sense and respond to extracellular guidance cues, and nucleokinesis, where interneurons propel their cell bodies forward in the selected direction of migration (Ang et al., 2003; Bellion et al., 2005; Moya and Valdeolmillos, 2004; Nadarajah et al., 2003; Polleux et al., 2002). Moreover, proper positioning and signaling from two subcellular organelles, the centrosome and primary cilium, have been implicated in the guided migration of cortical interneurons (Higginbotham et al., 2012; Luccardini et al., 2013; Luccardini et al., 2015; Yanagida et al., 2012). Failure to coordinate these cellular and subcellular events can alter cortical interneuron migration and impair the development of cortical circuitry, which may underlie severe neurological disorders such as autism spectrum disorder, schizophrenia, and epilepsy (Hildebrandt et al., 2011; Kato and Dobyns, 2005; Meechan et al., 2012; Volk et al., 2015). While progress has been made on elucidating the complex molecular mechanisms underlying nucleokinesis and leading process branching (Baudoin et al., 2012; Godin et al., 2012; Silva et al., 2018; Tsai and Gleeson, 2005), the intracellular signaling pathways that regulate these cellular mechanisms remain largely unknown.

The c-Jun NH_2_-terminal kinases (JNKs) are evolutionarily conserved members of the mitogen-activated protein kinase (MAPK) super-family (Chang and Karin, 2001; Davis, 2000). The JNK proteins are encoded by three genes, *Jnk1 (Mapk8), Jnk2 (Mapk9)*, and *Jnk3 (Mapk10).* JNKs phosphorylate numerous substrates in response to extracellular stimuli to mediate physiological processes including cellular proliferation, apoptosis, differentiation, and migration (Davis, 2000). Disruption to JNK signaling has been linked to aberrant migration of excitatory cortical neurons (Hirai et al., 2006; Wang et al., 2007; Westerlund et al., 2011; Yamasaki et al., 2011; Zhang et al., 2016) as well as cognitive disorders in humans (Kunde et al., 2013; McGuire et al., 2017). More recently, we found that JNK signaling controls the timing of interneuron entry into the cerebral cortex, as well as the formation and maintenance of tangential streams of cortical interneurons (Myers et al., 2020; Myers et al., 2014), but the role that JNK plays in the migratory properties of individual cortical interneurons has not been examined.

In the current study, we use a combination of pharmacological and genetic manipulations in an MGE explant cortical cell co-culture assay to demonstrate that interneurons have a requirement for JNK-signaling in the regulation of leading process branching and nucleokinesis. JNK-inhibited MGE interneurons dramatically slow their migration while displaying more variable speeds, and exhibit decreased migratory displacement. Concomitantly, JNK-inhibited interneurons display significant defects in leading process branching with decreased growth cone splitting frequency and interstitial side branch duration, as well as disrupted nucleokinesis and swelling dynamics. Similarly, genetic ablation of *Jnk* from MGE interneurons also results in leading process branching and nucleokinesis defects, suggesting interneurons have a cell-intrinsic requirement for JNK signaling during migration. In addition, we discovered a novel role for JNK signaling in the dynamic localization of the centrosome and primary cilium in migrating interneurons. Surprisingly, the centrosomes and the primary cilia of JNK-inhibited interneurons aberrantly localized to the cell body or trailing process, regardless of whether the leading process contained a swelling. These findings implicate the JNK pathway as a key intracellular mediator of leading process branching, nucleokinesis, and organelle dynamics in migrating MGE interneurons.

## RESULTS

### Pharmacological inhibition of JNK signaling disrupts MGE interneuron migration in vitro

c-Jun NH_2_-terminal kinase (JNK) signaling is required for the initial entry of cortical interneurons into the cortical rudiment and the tangential progression of interneurons in migratory streams (Myers et al., 2020; Myers et al., 2014). In the current study, we examined the role that JNK plays in the migratory dynamics of individual interneurons. To study interneuron migration at high spatial and temporal resolution, we performed live-cell confocal imaging of medial ganglionic eminence (MGE) explant cortical cell co-cultures. MGE explants from embryonic day 14.5 (E14.5) *Dlx5/6-Cre-IRES-EGFP (Dlx5/6-CIE)* positive embryos were cultured on top of a *Dlx5/6-CIE* negative (wild type, WT) monolayer of dissociated cortical cells for 24 hours (Fig. 1A). Cultures were treated with 20 μM SP600125, a pan JNK inhibitor (Bennett et al., 2001), or vehicle control and immediately imaged live for 12 hours (Fig. 1A). At the beginning of imaging (Time 0), the field of view was placed at the distal edge of interneuron outgrowth (Fig. 1B-C).

**Figure 1.**
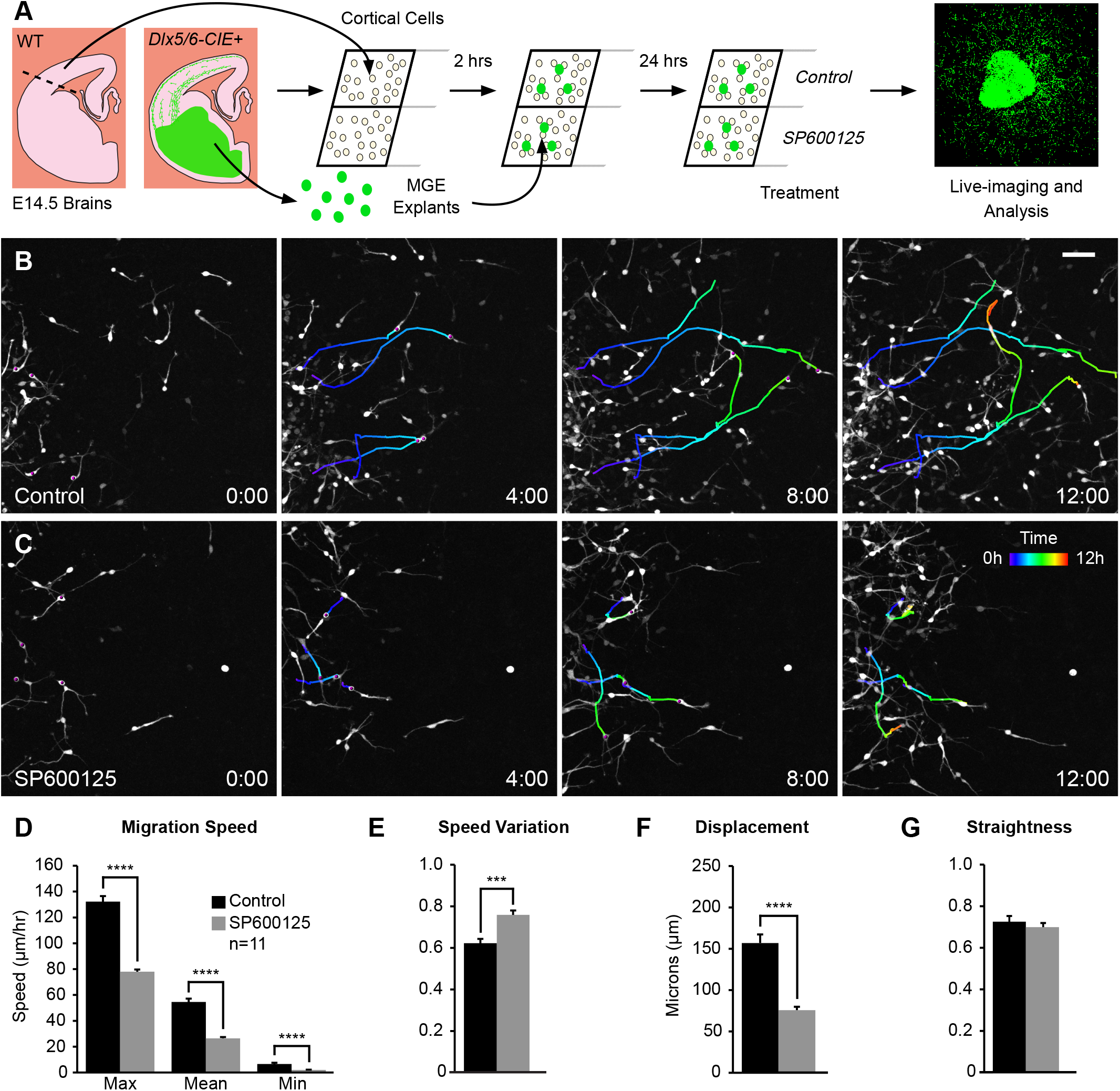
JNK signaling regulates the dynamic migratory properties of MGE interneurons. A. Schematic diagram of MGE explant cortical cell co-culture assay with pharmacological inhibition of JNK signaling. B-C. Individual cell tracks (pseudo-colored by time) from four interneurons in control (B) or 20μM SP600125 (C) treated cultures imaged live for 12 hours. D-G. Quantification of interneuron migratory properties revealed significant disruptions in migration speed (D), speed variation (E), and displacement (F), but not straightness (G) during JNK inhibition. For each condition, 10 cells were tracked from n = 11 movies (110 cells/condition) obtained over 4 experimental days. Data are mean±s.e.m. ****p<0.0001, ***p<0.001, Student’s *t*-test. Time in hours. Scale bar: 50 μm.

Many control interneurons migrated into the field of view by 12 hours of imaging (Fig. 1B; Movie 1), but SP600125-treated cells failed to progress through the frame and appeared to move slower (Fig. 1C; Movie 2). To assess potential differences in their migratory dynamics, we tracked individual cells in order to evaluate how JNK inhibition affects interneuron migration on a single cell level (representative cell tracks in Fig. 1B, C). The migratory speeds of JNK-inhibited interneurons were significantly slower than controls, including the maximum (values = mean±s.e.m.; control:132.28±4.25μm/hour; SP600125: 78.02±1.69μm/hour; p=1.68×10^-10^), mean (control: 54.62±2.54μm/hour; SP600125: 26.48±0.94μm/hour; p=1.68×10^-9^), and minimum (control: 6.64±0.91μm/hour; SP600125: 1.96±0.21 μm/hour; p=7.17×10^-5^) migratory speeds (Fig. 1D). While JNK-inhibited interneurons migrated slower, speed variation, which is the ratio of track standard deviation to track mean speed was significantly increased in SP600125-treated conditions (control: 0.62±0.02; SP600125 0.76±_-_0.02; p=0.00019; Fig. 1E). Due to the decrease in migratory speed, the normalized migratory displacement of SP600125-treated interneurons was also significantly reduced compared to control interneurons (control: 156.93±10.37μm; SP600125: 75.76±4.04 μm; p=4.73×10^-7^; Fig. 1F). Despite these changes in overall migratory dynamics, JNK-inhibited interneurons displayed no change in their migratory straightness (control: 0.71±0.03; SP600125: 0.68±0.02; p=0.45; Fig. 1G). Collectively, these data suggest that JNK inhibition alters the migratory behavior of MGE interneurons by reducing their migratory speed and the overall displacement of their migratory trajectories.

### JNK signaling regulates branching dynamics of migrating MGE interneurons

Migrating cortical interneurons repeatedly extend and retract leading process branches to sense extracellular guidance cues and establish a forward direction of movement (Bellion et al., 2005; Polleux et al., 2002; Yanagida et al., 2012). Leading process branching normally occurs through two mechanisms: growth cone splitting at the distal end of the leading process, and formation of interstitial side branches along the length of the leading process (Lysko et al., 2011; Martini et al., 2009).

To determine if JNK inhibition effected leading process morphology, we first measured the length of leading processes over time from live-imaged *Dlx5/6-CIE* positive MGE interneurons. Maximum (control: 84.96±4.45μm; SP600125: 85.14±4.02μm/hour; p=0.977), mean (control: 60.39±2.88μm; SP600125: 60.14±2.56μm; p=0.947), or minimum lengths (control: 37.40±2.47μm; SP600125: 37.81±2.72μm; p=0.912) of leading processes of SP600125-treated interneurons remained unchanged (Fig. 2C). However, when we analyzed the dynamic behavior of leading processes, significant differences were found between interneurons in control and SP600125-treated conditions (Fig. 2; Movies 3-4). In control conditions, migrating MGE interneurons show frequent initiation of new branches from growth cone splitting at the tip of their leading processes (Fig. 2A; Movie 3, Clip 1). In JNK-inhibited conditions, interneurons still underwent growth cone splitting, but the frequency appeared to be reduced (Fig. 2B; Movie 4, Clip1). When we measured the rate of growth cone splitting, JNK-inhibited interneurons had a statistically significant reduction compared to controls (control: 1.83±0.19 splits/hour; SP600125 1.15±0.20 splits/hour; p=0.02; Fig. 2D). In addition to branching from their growth cones, MGE interneurons extend and retract interstitial side branches from their leading processes. To determine whether JNK inhibition impacted the frequency and duration of interstitial branching, we measured the rate in which new side branches formed and determined the amount of time each newly generated branch was retained. Both control and SP600125-treated interneurons extended side branches at similar frequencies (control: 1.33±0.22 branches/hour; SP600125:1.37±0.19 branches/hour; p=0.91; Fig. 2 E-G; Movies 3-4, Clip 2). However, the duration of time in which *de novo* side branches persisted was significantly reduced in interneurons treated with JNK inhibitor (control: 28.77±2.53min; SP600125: 21.19±1.76min; p=0.02; Fig. 2H).

**Figure 2.**
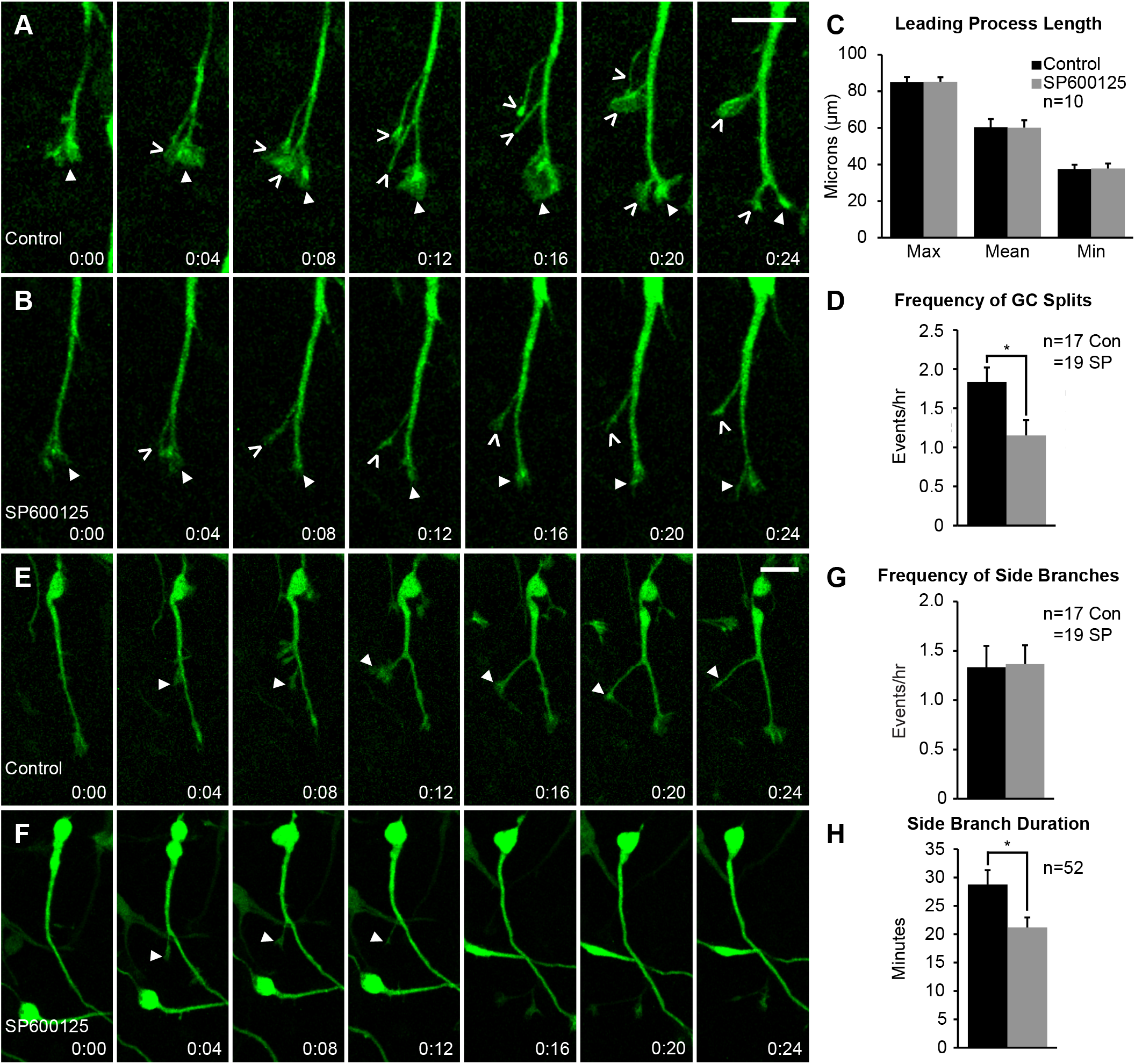
Migrating MGE interneurons require intact JNK signaling for proper leading process branching. A-B. Time series depicting growth cone (GC) splitting from control (A) or JNK-inhibited (B) MGE interneurons. Closed arrowhead = GC, open arrowhead = new GC branch. C. Quantification of leading process length. n=10 cells were measured from 8 movies/condition obtained over 4 experimental days. D. Quantification of GC splitting frequency. n=17 control cells from 8 movies and n=19 SP600125 cells from 10 movies were measured. E-F. Interstitial side branching from control (E) or JNK-inhibited (F) interneurons. Closed arrowhead = new side branch. G. Quantification of interstitial side branch frequency of control and SP600125 treated interneurons. n=17 control cells from 8 movies; n=19 SP600125 cells from 10 movies. H. Quantification of interstitial side branch duration in control and JNK-inhibited conditions. n=52 branches from 14 control cells and 18 SP600125 cells were measured from 10 movies/condition. All branching data were from movies obtained over 5 experimental days. Data are mean±s.e.m. *p<0.05, Student’s *t*-test. Time in minutes. Scale bar: 15 μm.

Here, we found that initiation of branching from growth cone splitting was significantly reduced during JNK inhibition. JNK-inhibited interneurons also formed side branches at similar rates, but these branches were shorter-lived than controls. Our data indicate that JNK influences branching dynamics of migratory MGE interneurons by regulating the rate of growth cone splitting, and by promoting the stability of newly formed side branches.

### Acute loss of JNK signaling impairs nucleokinesis and cytoplasmic swelling dynamics of migrating MGE interneurons

Since pharmacological inhibition of JNK signaling disrupted the overall migratory properties and leading process branching dynamics of MGE interneurons, we further examined the role for JNK in nucleokinesis, an obligate cell biological process in neuronal migration (Bellion et al., 2005; Yanagida et al., 2012). To closely examine the movement of interneuron cell bodies during migration, we imaged cultures at higher spatial and temporal resolution and analyzed the effect of JNK inhibition on nucleokinesis (Fig. 3). Time-lapse recordings show that under control conditions, a single cycle of nucleokinesis starts with the extension of a cytoplasmic swelling into the leading process and ends with the translocation of the cell body into the swelling (Fig. 3A; Movie 5, Clip 1). Although JNK-inhibited interneurons still engaged in nucleokinesis, the distance and kinetics of individual nucleokinesis events were disrupted (Fig. 3B; Fig. 3H; Movie 5, Clip 2). When we measured the mean distance that cell bodies advanced over time, JNK-inhibited interneurons translocated significantly shorter distances compared to control cells (control: 14.87±0.32μm; SP600125: 8.50±0.39μm; p=2.36×10^-10^; Fig. 3C). Thus, while cell bodies of JNK-inhibited interneurons still translocated forward into the leading process, the distance of their movement was reduced.

**Figure 3.**
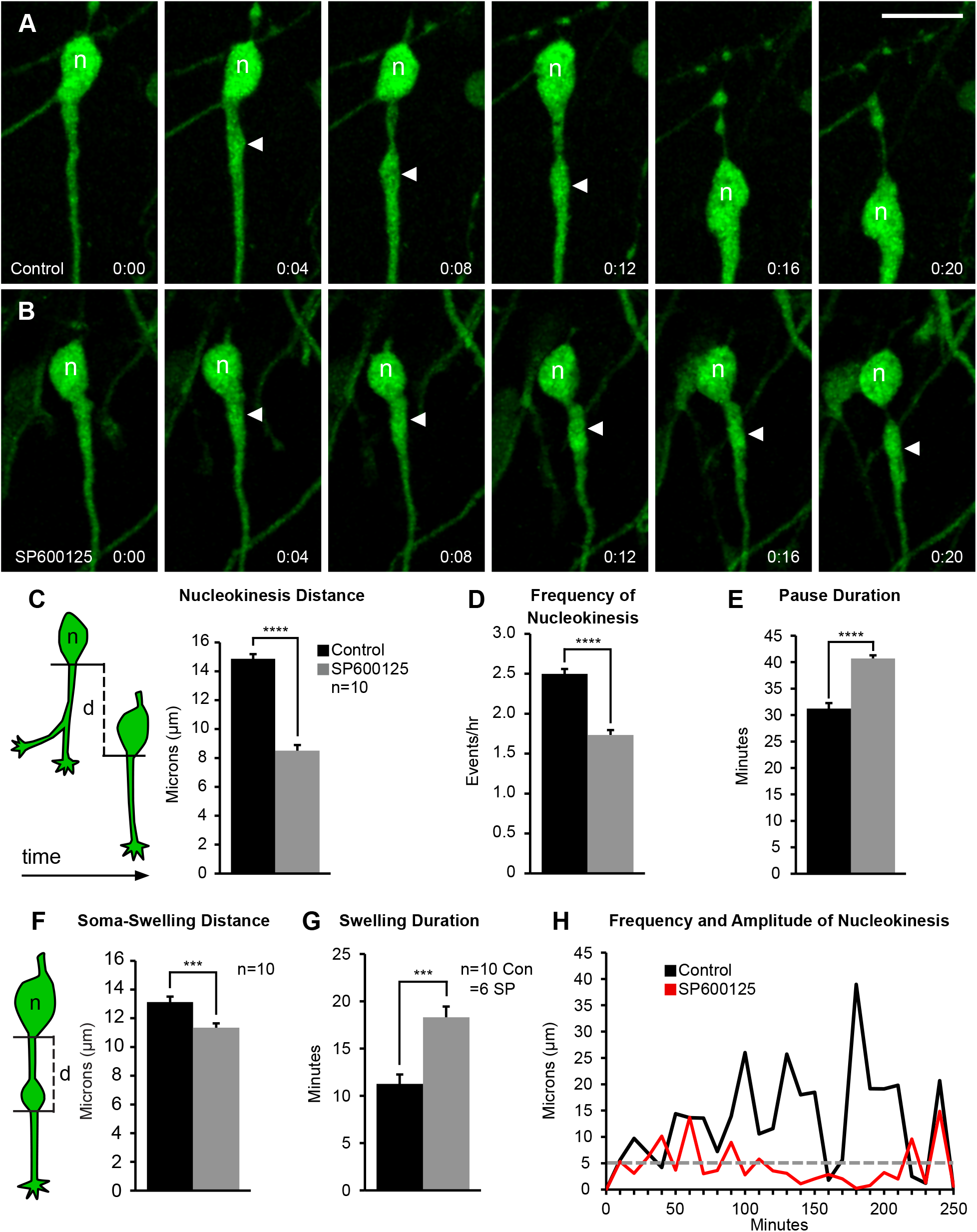
Pharmacological inhibition of JNK signaling impairs nucleokinesis in migrating MGE interneurons. A-B. Time series of a control (A) and SP600125-treated (B) interneuron undergoing a single cycle of nucleokinesis. Closed arrowhead = leading process swelling, n = nucleus. C-E. Cortical interneurons treated with JNK inhibitor have significantly shorter somal translocation distances (C), decreased frequency of nucleokinesis (D), and increased pause duration (E) compared to controls. C. Cartoon showing how the distance (d) that an interneuron cell body translocates over time was measured. In each condition, 50 cells were measured from n=10 movies obtained over 4 experimental days. F. Cartoon showing how the distance (d) that a swelling extends from a cell body was measured. JNK-inhibited cells display significantly decreased distance of swelling extension. G. Swelling duration is significantly increased in JNK-inhibited interneurons. 43 control cells were measured from n=10 control movies and 53 treated cells were measured from n=6 SP600125 movies, each obtained over 4 experimental days. H. Histogram showing nuclear translocation over time for a single cell in each condition. Distance traveled between two points is plotted and every movement above 5 μm (grey dashed line) is considered to be a nucleokinesis event. Data are mean±s.e.m. ****p<0.0001, ***p<0.001, **p<0.01, Student’s *t*-test. Time in minutes. Scale bar: 15 μm.

Since nucleokinesis is cyclical, with the cell extending a swelling, translocating its cell body, then pausing before repeating the process, we measured the rate of nucleokinesis in control and JNK-inhibited conditions. Upon treatment with SP600125, interneurons completed significantly fewer translocation events per hour (control: 2.50±0.06 events/hour; SP600125: 1.73±0.06 events/hour; p=1.92×10^-8^; Fig. 3D). Along with this, interneurons in JNK-inhibited cultures displayed longer pauses between the initiation of nucleokinesis events (control: 31.21±1.05min; SP600125: 40.71±0.58min; p=1.45×10^-7^; Fig. 3E). Because nuclear translocation is preceded by swelling extension, we measured the average distance from the soma to the swelling before translocation and found that SP600125-treated interneurons did not extend cytoplasmic swellings as far as controls (control: 13.13±0.38μm; SP600125: 11. 34±0.30μm; p=0.002; Fig. 3F). Since JNK-inhibited interneurons paused for longer periods of time, we asked if this was strictly due to delayed nuclear propulsion towards the swelling, or if the dynamics of swelling extension were also affected. Interneurons treated with SP600125 displayed significantly longer lasting cytoplasmic swellings (control: 11.27±0.99min; SP600125: 18.31±1.33min; p=0.0005; Fig. 3G), indicating that swelling duration is concomitantly increased with pause duration. Finally, the frequency and amplitude of nuclear translocations that exceed a minimum distance of 5 microns was notably reduced when individual control and JNK-inhibited cells were compared (Fig. 3H).

Together, these data point to a role for JNK signaling in regulating the distance and kinetics of nucleokinesis in migrating MGE interneurons, which likely contributes to the decrease in migratory speed and displacement that occurs during JNK inhibition.

### Complete genetic loss of JNK impairs nucleokinesis and leading process branching of migrating MGE interneurons in vitro

Since acute pharmacological inhibition of JNK activity altered the dynamic behavior of migratory cortical interneurons, we next asked whether genetic removal of JNK function from MGE interneurons also impaired their migration. In order to genetically ablate all three JNK genes from interneurons, we used mice containing the *Dlx5/6-CIE* transgene to conditionally remove *Jnk1* from *Jnk2;Jnk3* double knockout embryos (*Dlx5/6-CIE;Jnk1^fl/fl^;Jnk2^-/-^;Jnk3^-/-^*). Using this conditional triple knockout *(cTKO)* model, we modified our assay to determine if MGE interneurons have an intrinsic genetic requirement for JNK in their migration. MGE explants from *Dlx5/6-CIE+* wild type (WT) and *cTKO* brains were cultured on a WT cortical feeder layer and imaged live (Fig. 4A). We tracked individual interneurons over time to assess the overall migratory properties of WT and *cTKO* interneurons (Fig. 4B,C). While there were no changes in migratory speed (Fig. 4D), *cTKO* interneurons exhibited greater variations in migratory speed compared to WT cells (WT: 0.54±0.01; *cTKO:* 0.59±0.02; p=0.02; Figure 4E). We also found that *cTKO* interneurons have shorter migratory displacements than WT interneurons (WT: 195.06±6.80 μm; *cTKO:* 165.99±12.49 μm; p=0.05; Fig. 4F). Additionally, the track straightness of *cTKO* interneurons was decreased (WT: 0.77±0.02; *cTKO:* 0.71±0.02; p=0.03; Figure 4G). The combination of increased speed variability and decreased migratory straightness explain why *cTKO* interneurons exhibited shorter migratory displacements. Together, these data indicate that *cTKO* interneurons have subtle yet statistically significant deficits in their overall migratory dynamics, similar to pharmacological inhibition of JNK.

**Figure 4.**
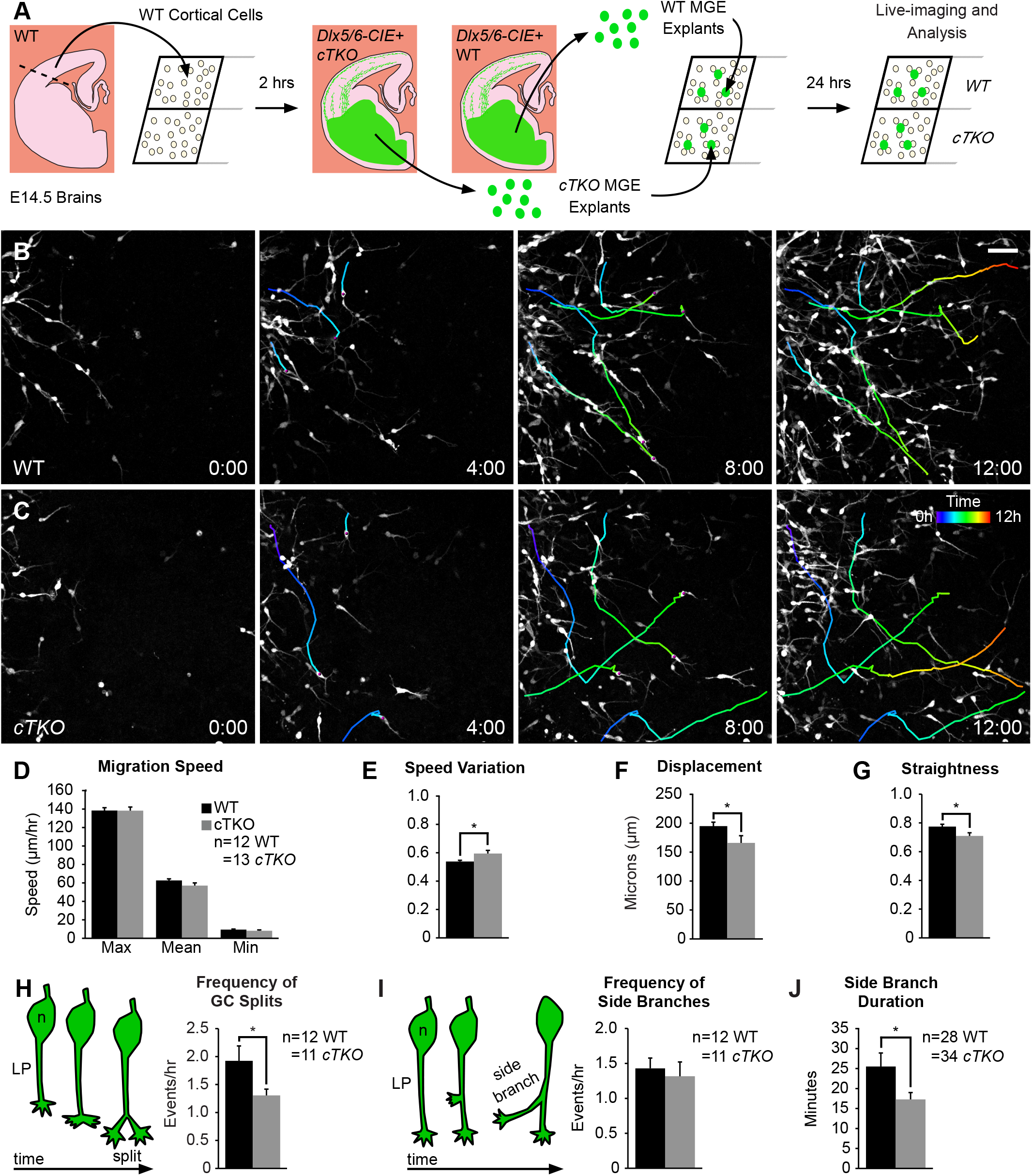
Genetic removal of JNK signaling impairs migratory properties and leading process dynamics of MGE interneurons. A. Diagram of MGE explant assay with *Dlx5/6-CIE+* wild-type (WT) or JNK conditional triple knockout (*cTKO*) explants cultured on WT cortical feeder-cells. B-C. Four individual cell tracks (pseudo-colored by time) from WT or *cTKO* interneurons imaged live for 12 hours. D-G. Quantification of migratory properties reveals significant disruptions in migratory speed, speed variation, displacement, and straightness between control and *cTKO* interneurons. 120 WT cells were measured from n=12 control movies and 130 *cTKO* cells were measured from n=13 *cTKO* movies, each obtained over 4 experimental days. H-I. *cTKO* interneurons have significantly decreased growth cone split frequency (H) without changes in interstitial side branch frequency (I). n=11 WT cells and n=12 *cTKO* cells measured from 6 movies/condition collected over 4 experimental days. J. Side branches from *cTKO* interneurons are significantly shorter-lived than controls. n=34 branches were measured from 10 WT cells and n=28 branches were measured from 10 *cTKO* cells recorded from 6 movies/condition obtained over 4 experimental days. Data are mean±s.e.m. *p<0.05, Student’s *t*-test. Time in hours. Scale bar: 50 μm.

To determine the genetic requirement for JNK signaling in branching, we analyzed leading process branching dynamics of *cTKO* and WT interneurons (Movies 6-7). *cTKO* interneurons displayed a significant reduction in the frequency of growth cone splitting compared to WT interneurons (WT: 1.92±0.18 splits/hour; *cTKO:* 1.30±0.11 splits/hour; p=0.04; Fig. 4H; Movie 6-7, Clip 1). In addition, genetic removal of JNK signaling from interneurons resulted in no change in side branch initiation (WT: 1.43±0.15; *cTKO*:1.32±0.20 branches/hour; p=0.66; Fig. 4I), but significant decreases in the duration that side branches persisted (WT: 25.51 ±3.39min; *cTKO*:17.29±1.71min; p=0.05; Fig. 4J; Movie 6-7, Clip 2). These data corroborate the findings from our pharmacological analyses and further suggest a key role for JNK signaling in controlling leading process branching dynamics.

Since we found alterations to overall migratory properties and branching dynamics, we next analyzed migrating *cTKO* interneurons for defects in nucleokinesis. Although *cTKO* interneurons engaged in nucleokinesis, the kinetics of nucleokinesis were significantly altered compared to WT interneurons (Fig. 5). The average distance *cTKO* cells traveled forward during nucleokinesis was significantly shorter compared to that of the WT cells (WT: 15.08±0.28μm; *cTKO:* 14.16±0.26μm; p=0.03; Fig. 5A-C). However, unlike during acute pharmacological inhibition of JNK signaling, *cTKO* interneurons displayed increased rates of nucleokinesis(Fig 5A, B; Movie 8). Genetic ablation of JNK signaling in migrating MGE interneurons resulted in increased frequency of translocation events (WT: 2.76±0.05 events/hour; *cTKO:* 3.22±0.11events/hour; p=0.002; Fig. 5D). While both WT and *cTKO* cells paused after the completion of a nucleokinesis event (after the cell body moves into the swelling), *cTKO* cells spent significantly less time pausing before they extended a new swelling (WT: 32.10±0.62min; *cTKO* 27.01 ±1.02min; p=0.0005; Fig. 5E). When we measured the duration of time that cytoplasmic swellings persisted, the swellings in *cTKO* interneurons were significantly shorter-lived (WT: 10.47±0.62 min; *cTKO:* 7.86±0.19min; p=0.003; Fig 5F). These data likely explain why we did not observe an overall change in migratory speeds between *cTKO* and WT interneurons. While *cTKO* interneurons are not migrating as far during each translocation event they are initiating nucleokinesis at a faster rate, thus moving at similar speeds compared to controls.

**Figure 5.**
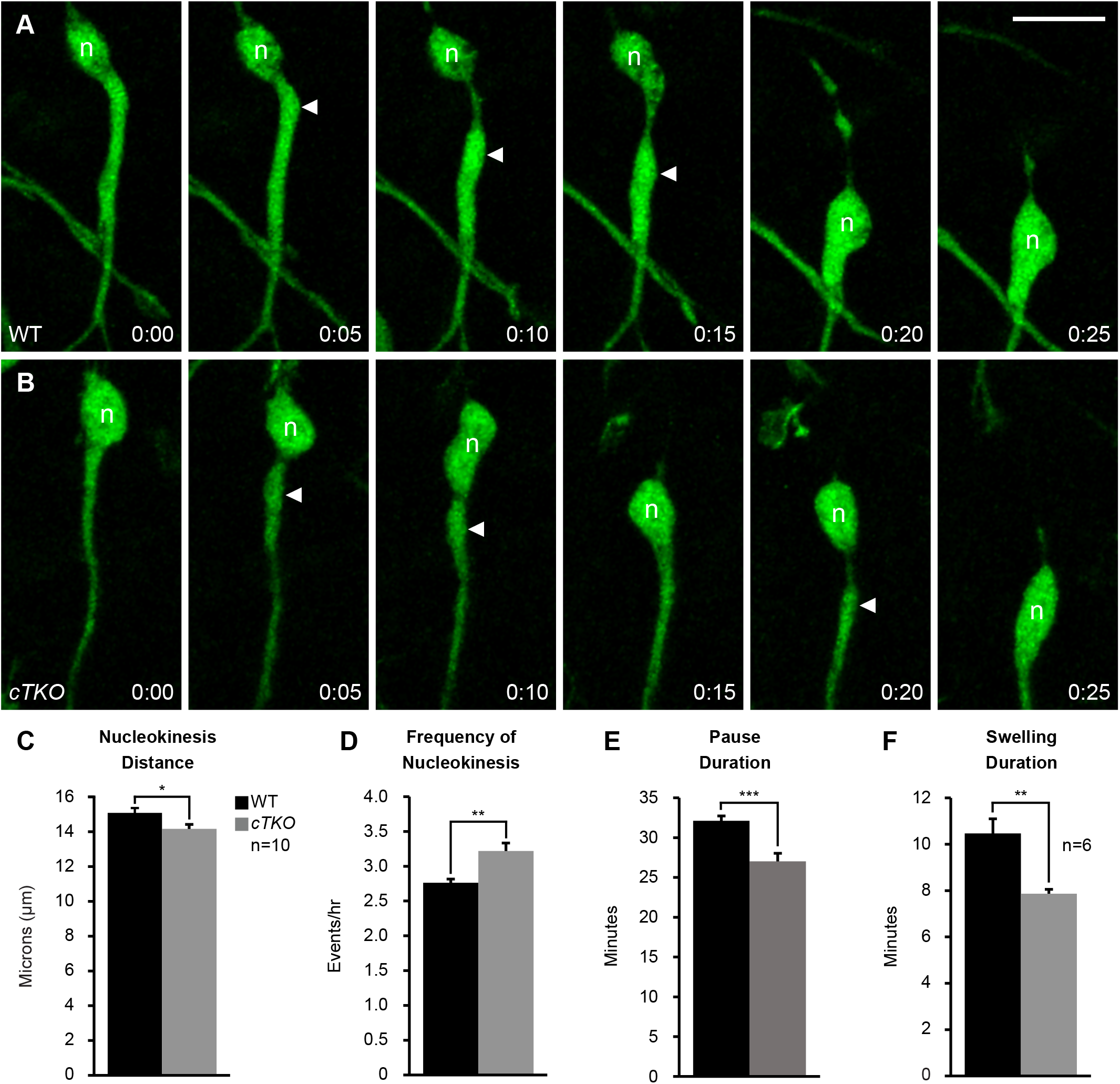
Genetic removal of *Jnk* disrupts nucleokinesis in migrating MGE interneurons. A. WT cortical interneuron undergoing a single nucleokinesis event. B. *cTKO* cortical interneuron completing two nucleokinesis events over the same interval of time. Closed arrowhead = leading process swelling, n = nucleus. C-E. *cTKO* interneurons have significantly decreased translocation distance (C), increased translocation frequency (D), and decreased pause duration (E) compared to WT interneurons. In each condition, 50 cells were measured from n=10 movies obtained over 4 experimental days. F. *cTKO* interneurons have decreased swelling duration compared to WT interneurons. 37 WT cells were measured from n=6 WT movies and 38 *cTKO* cells were measured from n=6 *cTKO* movies, each obtained over 4 experimental days. Data are mean±s.e.m. ***p<0.001, **p<0.01, *p<0.05, Student’s *t*-test. Time in minutes. Scale bar: 15 μm.

Collectively, our data suggest that genetic removal of *Jnk* alters the migratory behavior of MGE interneurons. While the phenotypes observed with conditional removal of *Jnk* from migrating interneurons was not identical to pharmacological inhibition of JNK signaling, our results indicate that interneurons require *Jnk* for correct leading process branching dynamics and nucleokinesis.

### Subcellular localization and dynamic behavior of the centrosome and primary cilia in migrating MGE interneurons depend on intact JNK-signaling

The cytoplasmic swelling emerges from the cell body during nucleokinesis and contains multiple subcellular organelles involved in the forward movement of cortical interneurons (Bellion et al., 2005; Martini and Valdeolmillos, 2010; Yanagida et al., 2012). One organelle involved in nucleokinesis is the centrosome, which translocates from the cell body into the swelling during nucleokinesis. The centrosome is tethered to the nucleus through a perinuclear cage of microtubules and acts to generate a forward pulling force on the nucleus during nucleokinesis (Bellion et al., 2005; Umeshima et al., 2007). Disruptions in centrosome motility and positioning are thought to underly nucleokinesis defects seen in other studies of neuronal migration (Luccardini et al., 2013; Luccardini et al., 2015; Silva et al., 2018; Solecki et al., 2009). Since we found significant defects in nucleokinesis in migrating MGE interneurons, we sought to determine if centrosome dynamics were also disrupted during JNK inhibition.

To visualize the centrosome and study the role of JNK signaling in centrosome dynamics in migrating MGE interneurons, we live-imaged *Dlx5/6-CIE+* cells expressing a red-fluorescent centrosome marker, Cetn2-mCherry (Fig. 6A). In control cells, the centrosome moved correctly into the cytoplasmic swelling (Fig 6B; Movie 9, Clip 1), with centrioles occasionally splitting between the soma and swelling preceding nucleokinesis (Fig. 6B, frames 0:00-0:10 minutes), as reported elsewhere (Bellion et al., 2005; Umeshima et al., 2007). Upon JNK-inhibition, the centrosome often maintained a position near the soma regardless of the presence of a swelling (Fig. 6C; Movie 9, Clip 2-3). Moreover, in many JNK-inhibited cells, the centrosome moved backwards into the trailing process, even when the cell body translocated forward (Fig. 6C; Movie 9, Clip 2-3). When we tracked the positioning of the centrosome over time, the centrosome of JNK-inhibited cells spent significantly more time in the trailing process and less time in the leading process (P=0.0001; Fig. 6D). Additionally, when a swelling was formed in front of the soma, the centrosome of JNK-inhibited cells spent significantly less time inside of the swelling than controls (control: 66.64±5.99%; SP600125: 16.08±5.52% of time; P=0.0001; Fig. 6E). When we measured the average maximal distance that the centrosome was displaced from the somal front, the centrosome of JNK-inhibited interneurons maintained a significantly closer position to the leading pole of the soma compared to controls (control: 9.93±0.99μm; SP600125: 6.73±0.88μm; p=0.03; Fig. 6F). This was not surprising since the soma-to-swelling distance in JNK-inhibited interneurons was decreased (Fig. 3E). However, when we compared the average maximal rearward distance between the centrosome and somal front, the centrosome of JNK-inhibited interneurons was significantly further behind that of controls(control: 9.40±0.77μm; SP600125: 19.75±1.94μm; p=1.48×10^-5^; Fig. 6F).

**Figure 6.**
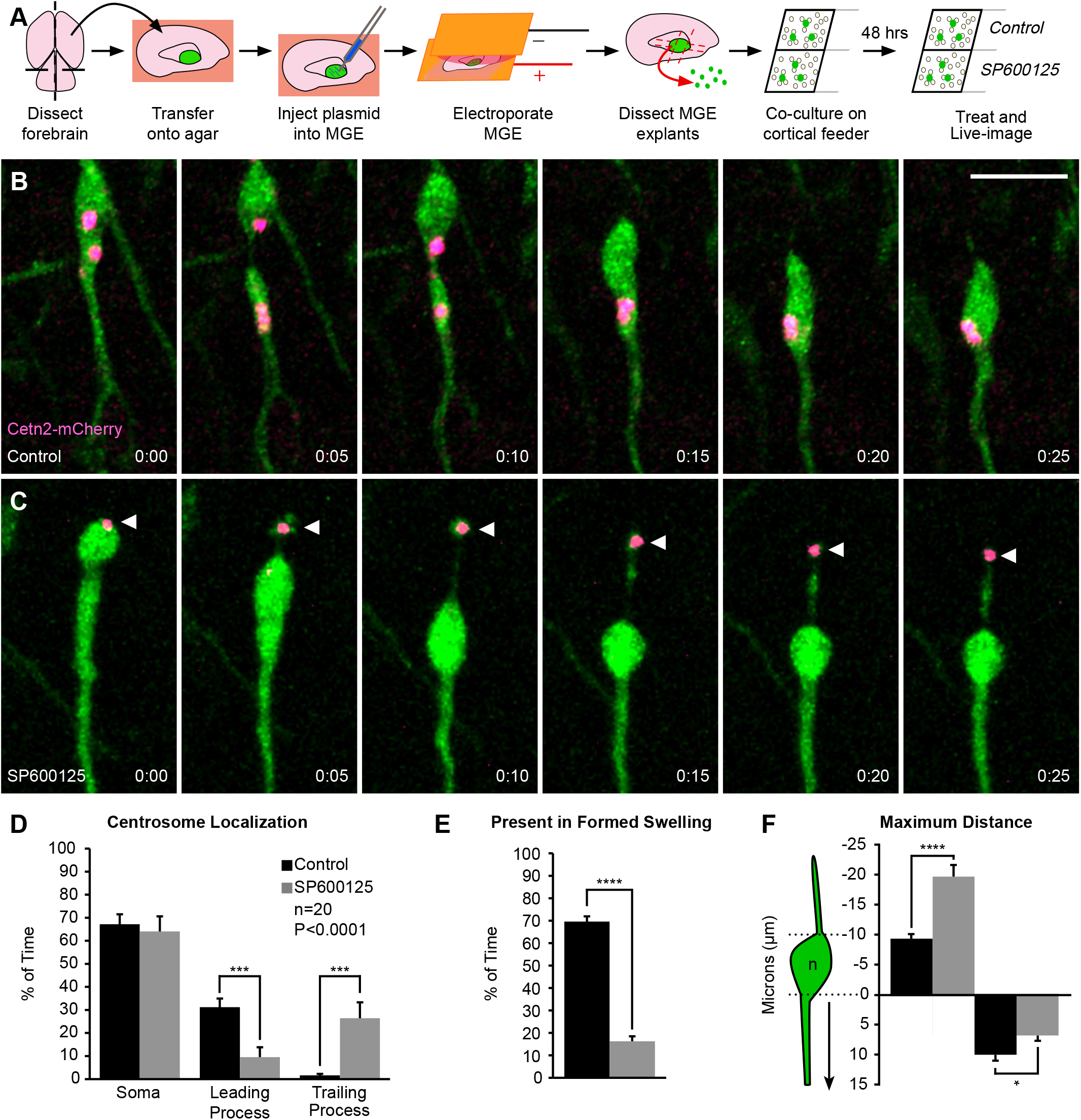
The subcellular localization of the centrosome is disrupted during JNK inhibition. A. Diagram depicting *ex vivo* electroporation of MGE tissue and subsequent culture of MGE explants on cortical feeder cells. B. An interneuron expressing a fluorescently tagged centrosome protein (Centrin2; Cetn2-mCherry) shows translocation of the centrosome into the cytoplasmic swelling prior to nucleokinesis in control conditions. C. A Cetn2-mCherry expressing interneuron treated with SP600125 shows aberrant rearward movement of the centrosome into the trailing process. Arrowhead = Cetn2-mCherry. D. Quantification of centrosome distribution over time (Two-way ANOVA: F(_2,114_) = 13.82; p<0.0001). Error bars represent mean±s.e.m., post-hoc by Fisher’s LSD ***p<0.001,**p<0.01, *p<0.05. E. Quantification of centrosome presence in a formed swelling over time (χ^2^ test; ****p<0.0001). F. Average maximum distance the centrosome traveled from the soma front (Student’s *t*-test; ****p<0.0001, ***p<0.001, **p<0.01, *p<0.05). In each condition, n=20 cells were measured from 11 movies obtained over 5 experimental days. Data are mean±s.e.m. Time in minutes. Scale bar: 15 μm.

Since we found defects in centrosome dynamics, we wanted to determine whether primary cilia, which normally extend from the mother centriole and house receptors important for the guided migration of cortical interneurons (Baudoin et al., 2012; Higginbotham et al., 2012), were also perturbed in interneurons following JNK-inhibition. In order to study the localization of cilia in migrating interneurons, we performed live-cell confocal imaging on *Dlx5/6-CIE+* MGE cells expressing Arl13b-tdTomato, a red-fluorescent cilia marker.

Almost identical to that of our centrosome analyses, we found significant alterations in the dynamic positioning of primary cilia in migrating MGE interneurons (Fig. 7). In control cells, the primary cilium moved into the cytoplasmic swelling before nuclear translocation (Fig. 7A; Movie 10, Clip 1). However, upon JNK inhibition, the cilium was frequently positioned in the soma and often moved into the trailing process as the cell body translocated forward (Fig. 7B; Movie 10, Clip 2-3). Overall, the cilia spent significantly more time in the cell soma and behind the cell in the trailing process, and significantly less time in the leading process of JNK-inhibited cells (P=0.0001; Fig. 7C). Additionally, the primary cilia in JNK-inhibited interneurons failed to spend as much time in formed cytoplasmic swellings as controls (control: 73.73±7.81% of time; SP600125: 33.85±8.20% of time; P=0.0001; Fig. 7D). When we measured the maximal distance behind the somal front, the cilia of JNK-inhibited interneurons were also positioned further behind the cell body than controls, matching our centrosome findings (control: 9.71±1.16μm; SP600125: 16.09±2.10μm; p=0.02; Fig. 7E). Taken together, these data highlight a novel role for JNK signaling in the dynamic movement and positioning of the centrosome and primary cilium in migrating MGE interneurons.

**Figure 7.**
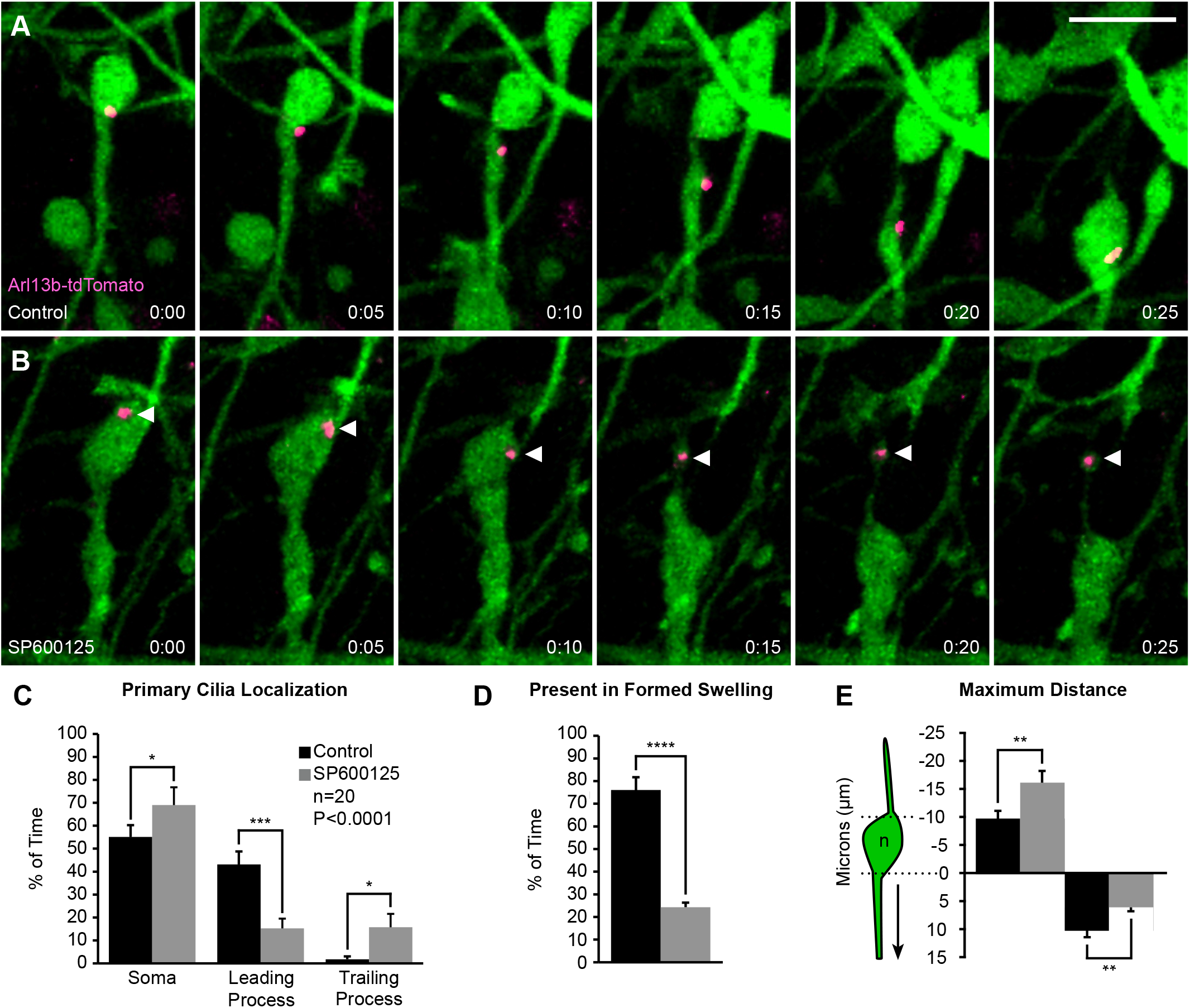
Primary cilium localization is disrupted during JNK inhibition. A. An interneuron expressing a fluorescently tagged primary ciliary marker (Arl13b-tdTomato) shows translocation of the primary cilium into the cytoplasmic swelling prior to nucleokinesis in control conditions. B. An interneuron expressing Arl13b-tdTomato shows aberrant rearward movement of the primary cilium into the trailing process when treated with SP600125. Arrowhead = Arl13b-tdTomato. C. Quantification of primary cilium distribution over time (Two-way ANOVA: F(_2,114_) = 12.13; p<0.0001). Error bars represent mean±s.e.m., post-hoc by Fisher’s LSD ***p<0.001,**p<0.01, *p<0.05. D. Quantification of primary cilium presence in a formed swelling over time (χ^2^ test; ****p<0.0001). E. Average maximum distance the primary cilium traveled from the soma front (Student’s *t*-test; **p<0.01). In each condition, n=20 cells were measured from 15 movies obtained over 6 experimental days. Data are mean±s.e.m. Time in minutes. Scale bar: 15 μm.

## DISCUSSION

In the present study, we demonstrated that migrating MGE interneurons rely on the JNK signaling pathway to properly undergo leading process branching and nucleokinesis. Pharmacological inhibition of JNK signaling in an *in vitro* assay resulted in reduced migratory speed and displacement with an increase in speed variation of migrating interneurons. Concomitant with these alterations in migratory properties, JNK-inhibited interneurons displayed decreased initiation of branches arising from growth cone tips, decreased persistence of interstitial side branches, as well as shorter, less frequent nucleokinesis events. Using a conditional triple knockout (*cTKO*) mouse line to completely remove *Jnk* from MGE interneurons, *cTKO* interneurons had decreased migratory displacement without reductions in overall migratory speed, apparently resulting from migratory trajectories that had more variable speeds and reduced track straightness compared to controls. Moreover, *cTKO* interneurons displayed significant defects in leading process branching and nucleokinesis. Similar to pharmacological manipulation, *cTKO* cells displayed shorter nuclear translocations, but unlike JNK-inhibited interneurons, *cTKO* interneurons completed nucleokinesis at faster rates relative to controls, which further explained why the overall migratory speed of *cTKO* interneurons was not impaired. These results indicate that MGE interneurons have a cell-intrinsic requirement in the coordination of leading process branching and nucleokinesis. Finally, we found a novel role of JNK signaling in regulating the dynamic positioning of two organelles involved in nucleokinesis: the centrosome and primary cilium. Centrosomes and primary cilia failed to properly translocate into a leading process swelling and spent significantly more time mislocalized to the trailing process of JNK-inhibited interneurons. Together, these results suggest that JNK signaling is required to maintain the cellular kinetics underlying MGE interneuron migration.

### Cytoskeletal regulation during leading process branching and nucleokinesis of migrating interneurons

Leading process branching and nucleokinesis—the two main features of guided interneuron migration—rely on the coordination of actomyosin and microtubule-based cytoskeletal networks. Leading process branches initially form through membrane protrusions containing a F-actin meshwork, which are then stabilized by microtubules to allow for the emergence of the nascent branch (Lysko et al., 2014; Martini et al., 2009; Peyre et al., 2015; Spillane et al., 2011). Nucleokinesis is thought to be mediated through the combination of the forward pulling forces from microtubules at the front of the cell and pushing forces from actomyosin contraction at the rear (Bellion et al., 2005; Martini and Valdeolmillos, 2010; Martini et al., 2009). While mechanisms underlying these processes are still under investigation, several molecular mediators of microtubule and actin dynamics in migrating interneurons have emerged, and interestingly, have been linked to JNK signaling in other cells.

For instance, p27^kip1^, a microtubule associated protein, coordinates both actomyosin contraction and microtubule organization to control leading process branching and nucleokinesis in migrating interneurons (Godin et al., 2012). Conditional deletion of p27^kip1^ from post-mitotic interneurons resulted in slower migratory speed, increased frequency of nucleokinesis, and shorter distance of translocations. Similarly, *cTKO* interneurons had shorter translocation distances and increased rates of nucleokinesis. In addition, p27^kip1^ knockout interneurons displayed shorter-lived side branches, similar to our findings with both pharmacological and genetic loss of JNK. JNK signaling was reported to regulate p27^kip1^ phosphorylation during cancer cell migration (Kim et al., 2012), suggesting a possible link between JNK signaling and this molecular mediator of cellular migration.

Another important regulator of nucleokinesis and leading process branching is the microtubule associated protein Doublecortin (Dcx; Friocourt et al., 2007; Kappeler et al., 2006), which is a downstream target of JNK signaling in neurons (Gdalyahu et al., 2004; Jin et al., 2010). Cortical interneurons lacking Dcx show a decreased duration of interstitial side branches, and significantly shorter nuclear translocation distances with no overall changes in migratory speed (Kappeler et al., 2006), similar to what we found in *cTKO* interneurons. Thus, it is possible that JNK signaling fine-tunes leading process branching and nucleokinesis in cortical interneurons by phosphorylating Dcx.

Recently, the role of the Elongator complex, specifically the enzymatic core Elp3, was found to control both leading process branching and nucleokinesis through the regulation of actomyosin activity (Tielens et al., 2016). MGE interneurons devoid of Elp3 displayed nucleokinesis and leading process branching defects strikingly similar to our pharmacological results, including decreased migratory speed, translocation frequency, nucleokinesis amplitude, and frequency of growth cone splitting (Tielens et al., 2016). Moreover, the Elongator complex was found to potentiate JNK activity during cellular stress in HeLa and HEK293 cells (Holmberg et al., 2002; Kojic and Wainwright, 2016). This suggests that the Elongator complex may potentiate the activity of JNK to phosphorylate effector proteins required for proper migration of interneurons. While the exact mechanisms underlying how cytoskeletal modulators interact to control the guided migration of cortical interneurons remain to be determined, JNK may be a key signaling node required to coordinate these cellular behaviors.

### Position and function of the centrosome and primary cilium during cortical interneuron migration

During the migration cycle of cortical interneurons, a cytoplasmic swelling containing two interconnected organelles, the centrosome and primary cilium, extends ahead of the soma into the leading process (Bellion et al., 2005; Tsai and Gleeson, 2005). Disruptions to the movement, positioning, and function of these organelles are often found in interneurons with migratory deficits (Baudoin et al., 2012; Higginbotham et al., 2012; Luccardini et al., 2013; Matsumoto et al., 2019; Nakamuta et al., 2017).

Migratory olfactory bulb interneurons require DOCK7, a member of the DOCK180 family of atypical Rac/Cdc42 guanine nucleotide exchange factors, for migration (Nakamuta et al., 2017). Knockdown of DOCK7 led to unstable movement of the centrosome from the swelling back into the cell body (Nakamuta et al., 2017), which was attributed to slower migration of olfactory bulb interneurons devoid of DOCK7. We observed similar migratory deficits and disrupted centrosome positioning in MGE interneurons treated with JNK inhibitor. Interestingly, knockdown of DOCK7 was previously shown to reduce JNK phosphorylation during Schwann cell development and migration (Yamauchi et al., 2008; Yamauchi et al., 2011).

Furthermore, inactivation of the cell adhesion molecule N-cadherin from MGE interneurons leads to mislocalization of the centrosome to the rear of the cell body (Luccardini et al., 2013). JNK-inhibition not only impeded the forward progression of centrosomes into the swelling, but also led to their unobstructed movement into the trailing process. Interestingly, JNK-inhibition has been reported to decrease N-cadherin levels and cellular migration of myofibroblasts (De Wever et al., 2004), which suggests a potential role for JNK signaling in the regulation of N-cadherin during migration. While mechanisms that control the positioning of the centrosome in migrating neurons remain to be explored, JNK signaling may help synchronize the activity of cell adhesion molecules, cytoskeletal proteins, and cytoplasmic machinery that are critically involved in centrosome motility.

Finally, disruptions to ciliary proteins including Arl13b, Kif3a, and IFT88 or to the sonic hedgehog (Shh) signal transduction pathway all result in cortical interneuron migratory deficits (Baudoin et al., 2012; Higginbotham et al., 2012). Conditional deletion of Arl13b disrupts the formation of the primary cilium from the centrosome and the localization/transport of key receptors known to be critical for interneuron migration, including C-X-C motif chemokine receptor 4 (Cxcr4), neuregulin-1 receptor (ErbB4), and the Serotonin Receptor 6 (5-Htr6) (Higginbotham et al., 2012; Riccio et al., 2009; Wang et al., 2011). Dominant negative knockdown of Kif3a, a molecular motor required for cilium-specific Shh signal transduction, results in rearward movement of the centrosome of migrating olfactory bulb interneurons (Matsumoto et al., 2019), suggesting that functional primary cilia are necessary for the proper localization of the centrosome-cilium complex. Thus, cortical interneurons may require the function of signal transduction machinery inside the primary cilium for the centrosome-cilium complex to localize correctly, and to sense and respond to environmental guidance cues that promote directed migration of interneurons. Additionally, cortical interneurons lacking Arl13b exhibited leading process branching defects, suggesting that the primary cilium may have cytoskeletal functions along with its role in transduction of guidance signals (Higginbotham et al., 2012). Here, we provided evidence that JNK signaling is required for the proper positioning of the primary cilium during MGE interneuron migration. Future studies are needed to determine whether inhibition of JNK signaling impairs the localization of centrosome and cilia by disrupting the function of ciliary proteins such as Kif3a, and whether mislocalized cilia can compromise the guided migration of cortical interneurons *in vivo.*

### Cellular influences of JNK signaling during cortical interneuron migration

Our work here has shown that the proper cellular mechanics of MGE interneuron migration depend on the JNK signaling pathway. Loss of JNK function disrupted leading process branching and nucleokinesis of MGE interneurons and led to significant alterations of their migratory properties. The requirement of JNK in interneuron migration could be multifactorial, however, and regulate interneuron migration through intrinsic mechanisms, extrinsic mechanisms, or both. Since SP600125 treatment inhibits JNK function in all cells of the MGE explant cortical cell co-culture assay, we cannot exclude the possibility that JNK inhibition disrupts cell-cell interactions between interneurons and the cortical feeder cells on which they are grown. To determine whether migrating MGE interneurons have a cell-autonomous requirement for JNK signaling, we genetically removed *Jnk* from interneurons and cultured them on WT cortical cells. Although we found migratory deficits in *cTKO* interneurons that were indicative of an intrinsic function for JNK, the deficits we uncovered were somewhat distinct from pharmacological experiments, suggesting that there may be additive effects when JNK is simultaneously removed from both populations of cells. Both pharmacological inhibition and genetic removal of *Jnk* resulted in consistent leading process branching phenotypes with decreased growth cone splitting and short-lived interstitial side branches. However, when we analyzed nucleokinesis, the kinetics of movement were opposite: JNK-inhibited cells completed nucleokinesis at slower rates, whereas *cTKO* cells completed at faster rates. These data imply that cortical interneuron migration is dependent on both intrinsic and extrinsic requirements for JNK signaling, as suggested from recent *in vivo* and *ex vivo* experiments (Myers et al., 2020). While the exact mechanisms that cortical interneurons utilize to navigate their environment remain to be fully elucidated, we have found that JNK signaling exerts fine-tune control over cell biological processes required for proper interneuron migration.

### Conclusions

Using a combination of pharmacological and genetic approaches, we found a novel requirement for JNK signaling in MGE interneuron leading process branching and nucleokinesis. Our findings are also the first to implicate the JNK signaling pathway as a key intracellular regulator of the dynamic positioning of multiple subcellular organelles involved in interneuron migration. The exact molecular mechanisms controlling JNK signaling in interneuron migration remain to be determined. Therefore, identifying the upstream activators and downstream targets of JNK signaling will provide further insight into the role of JNK signaling in cortical development and disease.

## MATERIALS AND METHODS

### Animals

Animals were housed and cared for by the Office of Laboratory Animal Resources at West Virginia University (Morgantown, WV, USA). Timed-pregnant dams (day of vaginal plug = embryonic day 0.5) were euthanized by rapid cervical dislocation at embryonic day 14.5 (E14.5) and mouse embryos were immediately harvested for tissue culture. CF-1 (Charles River; Wilmington, MA, US) or C57BL/6J dams (Stock # 000664; The Jackson Laboratory; Bar Harbour, ME, USA) were crossed to hemizygous *Dlx5/6-Cre-IRES-EGFP (Dlx5/6-CIE;* Stenman et al., 2003) males maintained on a C57BL/6J background to achieve timed pregnancies at E14.5. To generate JNK triple knockout embryos at E14.5, *Jnk1^fl/fl^; Jnk2^-/-^; Jnk3^-/-^* dams were crossed to *Dlx5/6-CIE; Jnk1^fl/+^; Jnk2^-/-^; Jnk3^+/-^* males maintained on a C57BL/6J background. All animal procedures were performed in accordance to protocols approved by the Institutional Animal Care and Use Committee at West Virginia University.

### MGE explant cortical cell co-culture

8-well chamber coverslip slides (Thermo Fisher 155411) were coated with a solution of poly-L-lysine (Sigma P5899) and laminin (Sigma L2020) diluted in sterile water (Polleux and Ghosh, 2002), incubated overnight at 37°C with 5% CO_2_, and rinsed with sterile water prior to cell plating. E14.5 *Dlx5/6-CIE+* and *Dlx5/6-CIE-* embryos were sorted by GFP fluorescence and dissected in ice-cold complete Hank’s Balanced Salt Solution (cHBSS; Tucker et al., 2006). Cortices were dissected from the negative brains and pooled together for dissociation (Polleux and Ghosh, 2002). After dissociation, 250μL of cell suspension diluted to 1680cells/μL was added to each well and allowed to settle for 2 hours. MGE explants were dissected from GFP+ brains and plated on top of cortical cells. Cultures were grown for 24 hours before treatments and live imaging. Two E14.5 timed-pregnant dams were used for each genetic experiment. *Dlx5/6-CIE+* and *Dlx5/6-CIE-* embryos were obtained from a *Dlx5/6-Cre-IRES-EGFP* x C57BL/6J cross, while *cTKO* embryos were obtained by crossing a *Dlx5/6-CIE; Jnk1^fl/+^; Jnk2^-/-^; Jnk3^+/-^* male to a *Jnk1^fl/fl^; Jnk2^-/-^; Jnk3^-/-^* dam. MGE explants from *Dlx5/6-CIE+* WT and *cTKO* embryos were dissected and plated into separate wells containing a monolayer of *Dlx5/6-CIE-* WT cortical cells. Cultures were grown 24 hours prior to live imaging.

### Electroporations

Intact ventral forebrains were microdissected from *Dlx5/6-CIE+* embryos and placed on thin slices of 3% low-melting point agarose (Fisher BP165-25) in cHBSS. Agar slices containing ventral forebrain tissue were placed onto a positive genepaddles electrode (5×7mm; Harvard Apparatus Inc #45-0123; Holliston, MA, USA) from a BTX ECM 830 squarewave electroporation system under a stereo microscope. Endotoxin-free plasmid DNA (1-3 mg/ml) for Cetn2-mCherry and Arl13b-tdTomato (gift from Dr. Eva Anton) was injected into the MGE with a picospritzer (6ms/spritz; General Valve Picospritzer II), a negative genepaddles electrode (5×7mm; Harvard Apparatus Inc #45-0123) containing a droplet of cHBSS was lowered to the tissue, and electroporated (5 x 60mV/5ms pulse length/200ms interval pulses). Electroporated MGE explants were then dissected, plated as above, and grown for 48 hours before imaging.

### Live Imaging Experiments

Cultures were treated with pre-warmed 37°C serum-free media containing a 1:1000 dilution of DMSO for vehicle control or 20 μM SP600125 pan-JNK inhibitor (Enzo Life Sciences BML-EI305-0010; Farmingdale, NY, USA) and immediately transferred to a Zeiss 710 Confocal Microscope with stable environmental controls maintained at 37°C with 5% humidified CO_2_. Multi-position time-lapse z-series were acquired at 10-minute intervals over a 12-hour period with a 20X Plan-Apo objective (Zeiss; Oberkochen, Germany) for overall migration analysis, nucleokinesis distance, and swelling distance measurements. For measurements requiring higher temporal and spatial resolution, such as swelling duration, branch dynamics, and visualization of subcellular structures in electroporated cells, cultures were imaged using multiposition time-lapse z-series at 2-2.5 minute intervals over a 4-10 hour period with a 40X C-apochromat 1.2W M27 objective (Zeiss; Oberkochen, Germany).

### Analysis of Live Imaging

4D live imaging movies were analyzed using Imaris 9.5.1 (Bitplane; Zürich, Switzerland) software. Movies collected at 20X were evaluated in the first 12 h of each recording. Individual interneurons were tracked for a minimum of 4 h. Tracks were discontinued if a cell remained stationary for 60 contiguous minutes, or if the tracked cell could no longer be unambiguously identified. All tracks from each movie were averaged together for dynamic analyses. Cortical interneurons were tracked using the Spots feature of Imaris to capture migratory speed, distance, displacement, and track straightness data. Displacement was normalized to the minimum track length of 4 h. Data sets were acquired from a minimum of four experimental days with genetic experiments containing 5 conditional triple knockout (*cTKO*) embryos. Pharmacological swelling duration data was obtained from movies collected over 4 experimental days. Genetic swelling duration was obtained from 3 experimental days with 3 *cTKO* embryos. The minimum criteria for an interstitial side branch to be included in our analysis was as follows: the cell had to remain in frame for a minimum of 3 hours, an interstitial side branch had to persist for a minimum of 10 minutes, and the branch could not become the new leading process. Two-tailed unpaired Student’s *t* tests were used to determine statistical differences between groups.

For electroporation experiments, cultures were imaged at 40X and cells were selected for centrosome and cilia analyses under the following criteria: the cell remained in frame for a minimum of 1 hour, the cell displayed low to moderate expression levels of the construct (without additional expression of aggregated fluorescent protein), and the cell was discernable from surrounding cells. Centrosome and ciliary distance from the front of the cell body, and localization were manually tracked and recorded using Imaris software. Two-way Anova followed by Fisher’s LSD post-hoc analyses were performed to determine statistical differences for organelle distribution analyses (Prism Version 8 using GraphPad Software; San Diego, CA, USA). Statistical significances were determined by χ^2^ test for the presence of absence of organelles to a formed swelling over time (Prism Version 8 using GraphPad Software; San Diego, CA, USA). Two-tailed unpaired Student’s *t* tests were used to determine statistical differences between groups for distance measurements. Confocal micrographs were uniformly adjusted for levels, brightness, and contrast in Imaris for movie preparation, and Adobe Photoshop for figure images.

## ACKNOWLEDGEMENTS

We would like to thank Dr. Amanda Ammer and Dr. Karen Martin for their excellent microscopy support. Live-imaging experiments were performed in the West Virginia University (WVU) Imaging Facilities, which were supported by the WVU Cancer Institute, the WVU Health Science Center Office of Research and Graduate Education, and NIH grants P20RR016440, P30GM103488, P20GM121322, U54GM104942, P30GM103503, and P20GM103434.

## AUTHOR CONTRIBUTIONS

Conceptualization: S.E.S. and E.S.T.; Methodology: S.E.S. and E.S.T.; Formal analysis: S.E.S., and N.K.C.; Investigation: S.E.S.; Writing: S.E.S., and E.S.T.; Visualization: S.E.S.; Supervision: E.S.T.; Funding Acquisition: E.S.T.

## SYMBOLS AND ABBREVIATIONS

MGE: Medial ganglionic eminence
CGE: Caudal ganglionic eminence
JNK: c-Jun NH_2_-terminal kinase
Dcx: Doublecortin
Cetn2-mCherry: Centrin2-mCherry
*Dlx5/6-CIE*: *Dlx5/6-Cre-IRES-EGFP*
*cTKO*: conditional *Jnk* triple knockout
WT: Wild type
MAPK: mitogen-activated protein kinase
cHBSS: complete Hank’s Balanced Salt Solution
Shh: Sonic hedgehog
E14.5: Embryonic day 14.5
μ: Micro
Cxcr4: C-X-C motif chemokine receptor 4
ErbB4: erb-b2 receptor tyrosine kinase 4
5-Htr6: Serotonin receptor 6
s.e.m.: standard error of the mean

## REFERENCES

Ang, E. S., Haydar, T. F., Gluncic, V. and Rakic, P. (2003). Four-dimensional migratory coordinates of GABAergic interneurons in the developing mouse cortex. J Neurosci 23, 5805–15.

Baudoin, J. P., Viou, L., Launay, P. S., Luccardini, C., Espeso Gil, S., Kiyasova, V., Irinopoulou, T., Alvarez, C., Rio, J. P., Boudier, T. et al. (2012). Tangentially migrating neurons assemble a primary cilium that promotes their reorientation to the cortical plate. Neuron 76, 1108–22.

Bellion, A., Baudoin, J. P., Alvarez, C., Bornens, M. and Métin, C. (2005). Nucleokinesis in tangentially migrating neurons comprises two alternating phases: forward migration of the Golgi/centrosome associated with centrosome splitting and myosin contraction at the rear. J Neurosci 25, 5691–9.

Bennett, B. L., Sasaki, D. T., Murray, B. W., O’Leary, E. C., Sakata, S. T., Xu, W., Leisten, J. C., Motiwala, A., Pierce, S., Satoh, Y. et al. (2001). SP600125, an anthrapyrazolone inhibitor of Jun N-terminal kinase. Proc Natl Acad Sci U S A 98, 13681–6.

Chang, L. and Karin, M. (2001). Mammalian MAP kinase signalling cascades. Nature 410, 37–40.

Davis, R. J. (2000). Signal transduction by the JNK group of MAP kinases. Cell 103, 239–52.

De Wever, O., Westbroek, W., Verloes, A., Bloemen, N., Bracke, M., Gespach, C., Bruyneel, E. and Mareel, M. (2004). Critical role of N-cadherin in myofibroblast invasion and migration in vitro stimulated by colon-cancer-cell-derived TGF-beta or wounding. J Cell Sci 117, 4691–703.

Friocourt, G., Liu, J. S., Antypa, M., Rakic, S., Walsh, C. A. and Parnavelas, J. G. (2007). Both doublecortin and doublecortin-like kinase play a role in cortical interneuron migration. J Neurosci 27, 3875–83.

Gdalyahu, A., Ghosh, I., Levy, T., Sapir, T., Sapoznik, S., Fishler, Y., Azoulai, D. and Reiner, O. (2004). DCX, a new mediator of the JNK pathway. EMBO J 23, 823–32.

Godin, J. D., Thomas, N., Laguesse, S., Malinouskaya, L., Close, P., Malaise, O., Purnelle, A., Raineteau, O., Campbell, K., Fero, M. et al. (2012). p27(Kip1) is a microtubule-associated protein that promotes microtubule polymerization during neuron migration. Dev Cell 23, 729–44.

Higginbotham, H., Eom, T. Y., Mariani, L. E., Bachleda, A., Hirt, J., Gukassyan, V., Cusack, C. L., Lai, C., Caspary, T. and Anton, E. S. (2012). Arl13b in primary cilia regulates the migration and placement of interneurons in the developing cerebral cortex. Dev Cell 23, 925–38.

Hildebrandt, F., Benzing, T. and Katsanis, N. (2011). Ciliopathies. N Engl J Med 364, 1533–43.

Hirai, S., Cui, D. F., Miyata, T., Ogawa, M., Kiyonari, H., Suda, Y., Aizawa, S., Banba, Y. and Ohno, S. (2006). The c-Jun N-terminal kinase activator dual leucine zipper kinase regulates axon growth and neuronal migration in the developing cerebral cortex. J Neurosci 26, 11992–2002.

Holmberg, C., Katz, S., Lerdrup, M., Herdegen, T., Jäättelä, M., Aronheim, A. and Kallunki, T. (2002). A novel specific role for I kappa B kinase complex-associated protein in cytosolic stress signaling. J Biol Chem 277, 31918–28.

Jin, J., Suzuki, H., Hirai, S., Mikoshiba, K. and Ohshima, T. (2010). JNK phosphorylates Ser332 of doublecortin and regulates its function in neurite extension and neuronal migration. Dev Neurobiol 70, 929–42.

Kappeler, C., Saillour, Y., Baudoin, J. P., Tuy, F. P., Alvarez, C., Houbron, C., Gaspar, P., Hamard, G., Chelly, J., Métin, C. et al. (2006). Branching and nucleokinesis defects in migrating interneurons derived from doublecortin knockout mice. Hum Mol Genet 15, 1387–400.

Kato, M. and Dobyns, W. B. (2005). X-linked lissencephaly with abnormal genitalia as a tangential migration disorder causing intractable epilepsy: proposal for a new term, “interneuronopathy”. J Child Neurol 20, 392–7.

Kim, H., Jung, O., Kang, M., Lee, M. S., Jeong, D., Ryu, J., Ko, Y., Choi, Y. J. and Lee, J. W. (2012). JNK signaling activity regulates cell-cell adhesions via TM4SF5-mediated p27(Kip1) phosphorylation. Cancer Lett 314, 198–205.

Kojic, M. and Wainwright, B. (2016). The Many Faces of Elongator in Neurodevelopment and Disease. Front Mol Neurosci 9, 115.

Kunde, S. A., Rademacher, N., Tzschach, A., Wiedersberg, E., Ullmann, R., Kalscheuer, V. M. and Shoichet, S. A. (2013). Characterisation of de novo MAPK10/JNK3 truncation mutations associated with cognitive disorders in two unrelated patients. Hum Genet 132, 461–71.

Luccardini, C., Hennekinne, L., Viou, L., Yanagida, M., Murakami, F., Kessaris, N., Ma, X., Adelstein, R. S., Mège, R. M. and Métin, C. (2013). N-cadherin sustains motility and polarity of future cortical interneurons during tangential migration. J Neurosci 33, 18149–60.

Luccardini, C., Leclech, C., Viou, L., Rio, J. P. and Métin, C. (2015). Cortical interneurons migrating on a pure substrate of N-cadherin exhibit fast synchronous centrosomal and nuclear movements and reduced ciliogenesis. Front Cell Neurosci 9, 286.

Lysko, D. E., Putt, M. and Golden, J. A. (2011). SDF1 regulates leading process branching and speed of migrating interneurons. J Neurosci 31, 1739–45.

Lysko, D. E., Putt, M. and Golden, J. A. (2014). SDF1 reduces interneuron leading process branching through dual regulation of actin and microtubules. J Neurosci 34, 4941–62.

Martini, F. J. and Valdeolmillos, M. (2010). Actomyosin contraction at the cell rear drives nuclear translocation in migrating cortical interneurons. J Neurosci 30, 8660–70.

Martini, F. J., Valiente, M., López Bendito, G., Szabó, G., Moya, F., Valdeolmillos, M. and Marín, O. (2009). Biased selection of leading process branches mediates chemotaxis during tangential neuronal migration. Development 136, 41–50.

Matsumoto, M., Sawada, M., García-González, D., Herranz-Pérez, V., Ogino, T., Bang Nguyen, H., Quynh Thai, T., Narita, K., Kumamoto, N., Ugawa, S. et al. (2019). Dynamic Changes in Ultrastructure of the Primary Cilium in Migrating Neuroblasts in the Postnatal Brain. J Neurosci 39, 9967–9988.

McGuire, J. L., Depasquale, E. A., Funk, A. J., O’Donnovan, S. M., Hasselfeld, K., Marwaha, S., Hammond, J. H., Hartounian, V., Meador-Woodruff, J. H., Meller, J. et al. (2017). Abnormalities of signal transduction networks in chronic schizophrenia. NPJ Schizophr 3, 30.

Meechan, D. W., Tucker, E. S., Maynard, T. M. and LaMantia, A. S. (2012). Cxcr4 regulation of interneuron migration is disrupted in 22q11.2 deletion syndrome. Proc Natl Acad Sci U S A 109, 18601–6.

Miyoshi, G., Hjerling-Leffler, J., Karayannis, T., Sousa, V. H., Butt, S. J., Battiste, J., Johnson, J. E., Machold, R. P. and Fishell, G. (2010). Genetic fate mapping reveals that the caudal ganglionic eminence produces a large and diverse population of superficial cortical interneurons. J Neurosci 30, 1582–94.

Moya, F. and Valdeolmillos, M. (2004). Polarized increase of calcium and nucleokinesis in tangentially migrating neurons. Cereb Cortex 14, 610–8.

Myers, A. K., Cunningham, J. G., Smith, S. E., Snow, J. P., Smoot, C. A. and Tucker, E. S. (2020). JNK signaling is required for proper tangential migration and laminar allocation of cortical interneurons. Development 147, Doi: 10.1242/dev.180646.

Myers, A. K., Meechan, D. W., Adney, D. R. and Tucker, E. S. (2014). Cortical interneurons require Jnk1 to enter and navigate the developing cerebral cortex. J Neurosci 34, 7787–801.

Nadarajah, B., Alifragis, P., Wong, R. O. and Parnavelas, J. G. (2003). Neuronal migration in the developing cerebral cortex: observations based on real-time imaging. Cereb Cortex 13, 607–11.

Nakamuta, S., Yang, Y. T., Wang, C. L., Gallo, N. B., Yu, J. R., Tai, Y. and Van Aelst, L. (2017). Dual role for DOCK7 in tangential migration of interneuron precursors in the postnatal forebrain. J Cell Biol 216, 4313–4330.

Nery, S., Fishell, G. and Corbin, J. G. (2002). The caudal ganglionic eminence is a source of distinct cortical and subcortical cell populations. Nat Neurosci 5, 1279–87.

Peyre, E., Silva, C. G. and Nguyen, L. (2015). Crosstalk between intracellular and extracellular signals regulating interneuron production, migration and integration into the cortex. Front Cell Neurosci 9, 129.

Polleux, F. and Ghosh, A. (2002). The slice overlay assay: a versatile tool to study the influence of extracellular signals on neuronal development. Sci STKE 2002, pl9.

Polleux, F., Whitford, K. L., Dijkhuizen, P. A., Vitalis, T. and Ghosh, A. (2002). Control of cortical interneuron migration by neurotrophins and PI3-kinase signaling. Development 129, 3147–60.

Riccio, O., Potter, G., Walzer, C., Vallet, P., Szabó, G., Vutskits, L., Kiss, J. Z. and Dayer, A. G. (2009). Excess of serotonin affects embryonic interneuron migration through activation of the serotonin receptor 6. Mol Psychiatry 14, 280–90.

Silva, C. G., Peyre, E., Adhikari, M. H., Tielens, S., Tanco, S., Van Damme, P., Magno, L., Krusy, N., Agirman, G., Magiera, M. M. et al. (2018). Cell-Intrinsic Control of Interneuron Migration Drives Cortical Morphogenesis. Cell 172, 1063–1078.e19.

Solecki, D. J., Trivedi, N., Govek, E. E., Kerekes, R. A., Gleason, S. S. and Hatten, M. E. (2009). Myosin II motors and F-actin dynamics drive the coordinated movement of the centrosome and soma during CNS glial-guided neuronal migration. Neuron 63, 63–80.

Spillane, M., Ketschek, A., Jones, S. L., Korobova, F., Marsick, B., Lanier, L., Svitkina, T. and Gallo, G. (2011). The actin nucleating Arp2/3 complex contributes to the formation of axonal filopodia and branches through the regulation of actin patch precursors to filopodia. Dev Neurobiol 71, 747–58.

Stenman, J., Toresson, H. and Campbell, K. (2003). Identification of two distinct progenitor populations in the lateral ganglionic eminence: implications for striatal and olfactory bulb neurogenesis. J Neurosci 23, 167–74.

Tielens, S., Huysseune, S., Godin, J. D., Chariot, A., Malgrange, B. and Nguyen, L. (2016). Elongator controls cortical interneuron migration by regulating actomyosin dynamics. Cell Res 26, 1131–1148.

Tsai, L. H. and Gleeson, J. G. (2005). Nucleokinesis in neuronal migration. Neuron 46, 383–8.

Tucker, E. S., Polleux, F. and LaMantia, A. S. (2006). Position and time specify the migration of a pioneering population of olfactory bulb interneurons. Dev Biol 297, 387–401.

Umeshima, H., Hirano, T. and Kengaku, M. (2007). Microtubule-based nuclear movement occurs independently of centrosome positioning in migrating neurons. Proc Natl Acad Sci U S A 104, 16182–7.

Volk, D. W., Chitrapu, A., Edelson, J. R. and Lewis, D. A. (2015). Chemokine receptors and cortical interneuron dysfunction in schizophrenia. Schizophr Res 167, 12–7.

Wang, X., Nadarajah, B., Robinson, A. C., McColl, B. W., Jin, J. W., Dajas-Bailador, F., Boot-Handford, R. P. and Tournier, C. (2007). Targeted deletion of the mitogen-activated protein kinase kinase 4 gene in the nervous system causes severe brain developmental defects and premature death. Mol Cell Biol 27, 7935–46.

Wang, Y., Li, G., Stanco, A., Long, J. E., Crawford, D., Potter, G. B., Pleasure, S. J., Behrens, T. and Rubenstein, J. L. (2011). CXCR4 and CXCR7 have distinct functions in regulating interneuron migration. Neuron 69, 61–76.

Westerlund, N., Zdrojewska, J., Padzik, A., Komulainen, E., Björkblom, B., Rannikko, E., Tararuk, T., Garcia-Frigola, C., Sandholm, J., Nguyen, L. et al. (2011). Phosphorylation of SCG10/stathmin-2 determines multipolar stage exit and neuronal migration rate. Nat Neurosci 14, 305–13.

Wichterle, H., Garcia-Verdugo, J. M., Herrera, D. G. and Alvarez-Buylla, A. (1999). Young neurons from medial ganglionic eminence disperse in adult and embryonic brain. Nat Neurosci 2, 461–6.

Xu, Q., Cobos, I., De La Cruz, E., Rubenstein, J. L. and Anderson, S. A. (2004). Origins of cortical interneuron subtypes. J Neurosci 24, 2612–22.

Yamasaki, T., Kawasaki, H., Arakawa, S., Shimizu, K., Shimizu, S., Reiner, O., Okano, H., Nishina, S., Azuma, N., Penninger, J. M. et al. (2011). Stress-activated protein kinase MKK7 regulates axon elongation in the developing cerebral cortex. J Neurosci 31, 16872–83.

Yamauchi, J., Miyamoto, Y., Chan, J. R. and Tanoue, A. (2008). ErbB2 directly activates the exchange factor Dock7 to promote Schwann cell migration. J Cell Biol 181, 351–65.

Yamauchi, J., Miyamoto, Y., Hamasaki, H., Sanbe, A., Kusakawa, S., Nakamura, A., Tsumura, H., Maeda, M., Nemoto, N., Kawahara, K. et al. (2011). The atypical Guanine-nucleotide exchange factor, dock7, negatively regulates schwann cell differentiation and myelination. J Neurosci 31, 12579–92.

Yanagida, M., Miyoshi, R., Toyokuni, R., Zhu, Y. and Murakami, F. (2012). Dynamics of the leading process, nucleus, and Golgi apparatus of migrating cortical interneurons in living mouse embryos. Proc Natl Acad Sci U S A 109, 16737–42.

Zhang, F., Yu, J., Yang, T., Xu, D., Chi, Z., Xia, Y. and Xu, Z. (2016). A Novel c-Jun N-terminal Kinase (JNK) Signaling Complex Involved in Neuronal Migration during Brain Development. J Biol Chem 291, 11466–75.

